# Plasma lipidome dysregulation in frontotemporal dementia reveals shared, genotype-specific, and severity-linked alterations

**DOI:** 10.1101/2025.04.20.649464

**Authors:** Yohannes A. Ambaw, Peter A. Ljubenkov, Shubham Singh, Abdi Hamed, Sebastian Boland, Adam L. Boxer, Tobias C. Walther, Robert V. Farese

**Affiliations:** Cell Biology Program, Sloan Kettering Institute, New York, NY, USA; Department of Neurology, Memory and Aging Center, University of California San Francisco, San Francisco, CA, USA; Department of Molecular Metabolism, Harvard T.H. Chan School of Public Health, Boston, MA, USA; Peresent affiliation: Gene Therapy Department, Eli Lilly and Company, New York, NY, USA; Howard Hughes Medical Institute, New York, NY, USA

**Keywords:** Frontotemporal dementia, lipidomics, biomarker, *GRN*, *C9orf72*, *MAPT*, and sphingolipids

## Abstract

Developing new treatment strategies for frontotemporal dementia (FTD) and other forms of neurodegeneration requires biomarkers to monitor disease progression. Dysregulated brain lipid metabolism, particularly sphingolipids enriched in the nervous system, is a key feature of neurodegeneration. However, plasma lipids have been investigated less for their potential as biomarkers than brain imaging and serum proteins. Here we examined the plasma lipidomes of a cohort of heterozygous carriers of gene variants associated with autosomal dominant familial FTD (including *GRN*, *C9orf72*, and *MAPT* loci), comparing them with aged-matched controls. In general, FTD subjects exhibited increases in plasma levels of specific species of gangliosides (GM3(d18:1_16:0), GM3(d18:1_18:0), and GM3(d18:1_24:1)) and ceramide (Cer(d18:1_23:0)) and selected polyunsaturated triacylglycerols (TG). Other species of ceramides (Cer(d18:0_22:0)), phosphatidylethanolamine (PE(18:0_24:0)), and sphingomyelin (SM(38:0)) were reduced in plasma of FTD subjects. Levels of glucosylsphingosine (GlcSph(d18:1)) were elevated specifically in *GRN* carriers, SM(34:1) was reduced in *C9orf72* carriers, and TG(16:0_18:1_20:3)) were decreased in *MAPT* variant carriers. Notably, the ganglioside GM3(d18:1_16:0) was consistently elevated across all FTD genetic subtypes. Furthermore, the levels of these lipids correlated with disease severity in FTD patients. Our findings suggest that specific plasma lipid changes, notably several sphingolipids, may be useful biomarkers for FTD disease or progression.

## Introduction

Frontotemporal dementia (FTD) is a progressive neurodegenerative disease that typically presents with disabling changes in behavior or language under age 65 [1–3]. The neuropathological correlate of FTD, frontotemporal lobar degeneration (FTLD), most commonly involves neuronal mislocalization of transactive response DNA-binding protein 43 (TDP-43) or aggregation of tau isoforms [4, 5]. FTD is commonly associated with a strong family history of neurodegenerative disease, and about a quarter of FTD can be attributed to an autosomal dominant familial form of FTLD, mostly typically hexanucleotide expansion of chromosome 9 open reading frame 72 (*C9orf72*), progranulin gene (*GRN*) haploinsufficiency, and pathogenic variants of microtubule associated protein tau (*MAPT*) [6–8].

There are currently no effective treatments for FTD, and versatile fluid biomarkers for diagnosis and disease progression are necessary to investigate whether future therapeutic candidates are able to target specific pathogenic mechanisms. To date relatively nonspecific measures of neuronal injury, such as plasma levels of neurofilament light chain (NfL) [9, 10] or glial function, such as glial fibrillary acidic protein (GFAP) [11], and other proteomic biomarkers emerged as potential candidate pharmacodynamic biomarkers for FTD clinical trials, while lipidomic biomarkers have been less investigated. Yet, the brain is made up predominantly of lipids, constituting ∼60% of its dry mass [12, 13], and compared with many peripheral tissues, is highly enriched for sphingolipids, including sphingomyelin and glycosphingolipids. Recent studies of FTD subjects show alterations in sphingolipids in samples of diseased brains. Moreover, post-mortem brain samples from subjects with FTD from *GRN* haploinsufficiency contain elevated levels of gangliosides and are deficient in lysosomal bis(monoacylglycero)phosphate (BMP) lipids [4, 14]. Gangliosides are complex glycosphingolipids containing sialic acid, and BMP is required for their efficient degradation [4, 14]. Similar changes in gangliosides and BMP were found in mice or human cultured cell lines with in progranulin deficiency [15, 16]. Additionally, the levels of some sphingolipid degradation products, such as glucosylsphingosine (GlcSph), are elevated in plasma of subjects with FTD-*GRN* [17, 18]. Thus, plasma lipidomics may provide additional candidate biomarkers of FTD biology to support future clinical therapeutic development To systematically test whether plasma lipids may serve as useful biomarkers for FTD, we investigated the plasma lipidomes of individuals with *GRN*, *C9orf72*, and *MAPT* variants, comparing the results with healthy, age-matched controls. Familial forms of FTD were selected to enable comparisons between cohorts with more predictable associations with specific FTD subtypes and underlying mechanisms of pathogenicity than in sporadic FTD. Our measurements included high-resolution mass spectrometry–based lipidomics that detect a large variety of glycerolipids, along with a specific, separate analysis for more amphipathic glycosphingolipids. We also assessed the relationship between specific plasma lipid levels and disease severity. Our results provide new insights into plasma lipid biomarkers that may have potential utility for monitoring FTD disease progression.

## Results

### Characteristics of the study populations

We analyzed 130 plasma samples from patients with various forms of FTD, including those with mutations of *GRN*, *C9orf72*, and *MAPT*, as well as from control subjects (Figure 1). Table 1 lists the clinical and demographic characteristics of the participants in this study. The mean age was 49.2 ± 11.7 years for the control group, 56.1 ± 11.1 years for the FTD-*C9orf72* group, 61.4 ± 10.7 years for the FTD-*GRN* group, and 54.5 ± 7.8 years for the FTD-*MAPT* group. Most participants (98.5%) were of Caucasian origin, and symptomatic FTD cases exhibited mixed clinical phenotypes with respect to behavioral, aphasic variants, and other phenotypes. A fraction of the symptomatic mutation carriers had the behavioral variant of FTD (bvFTD; 46% in FTD-*GRN*, 92% in FTD-*C9orf72*, 100% in FTD-*MAPT*), with a lower percentage of patients showing primary progressive aphasia (PPA; 25% in FTD-*GRN*, 7.2% in FTD-*C9orf72,* 0% in FTD-*MAPT*).

**Figure 1.**
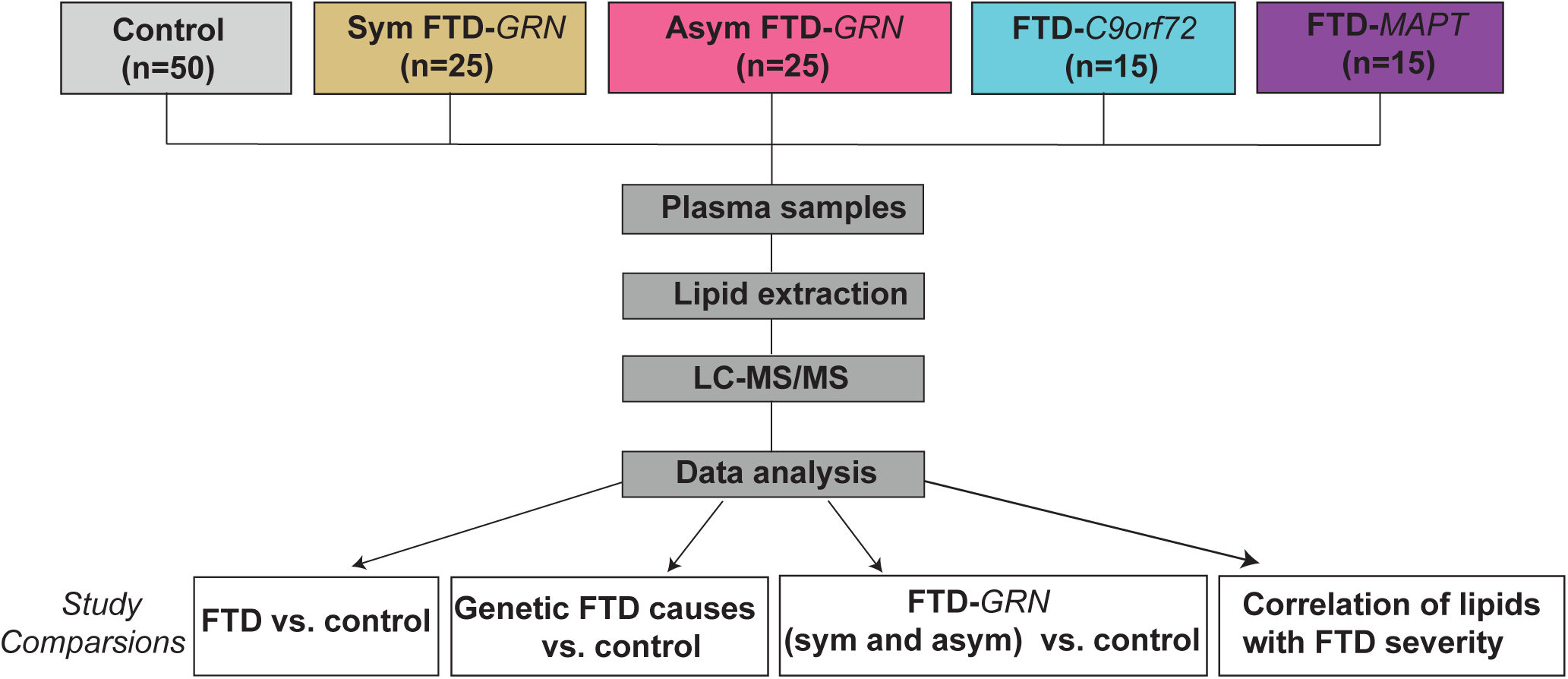
**Flow diagram of study design and workflow**, including study groups, experimental steps, and study conditions.

**Table 1:** Demographics and clinical characteristics for biofluid samples.

|  | Control | FTD |  |  |  |
| --- | --- | --- | --- | --- | --- |
|  |  | FTD-GRN sym | FTD-GRN asym | FTD-C9orf72 | FTD-MAPT |
| **Age** |  |  |  |  |  |
| Mean (SD) | 49.2 (11.7) | 65.4 (10.7) | 51.3 (16.9) | 56.1 (11.1) | 54.5 (7.8) |
| **Sex** |  |  |  |  |  |
| Male (n) | 26 | 12 | 11 | 6 | 7 |
| Female (n) | 24 | 13 | 14 | 9 | 7 |
| **CDR® plus NACC**<br>**FTLD Sum of Boxes**<br>**Score at First Visit** |  |  |  |  |  |
| Mean (SD) | 0 (0) | 10.5 (7.4) | 0.36(1.6) | 8.03(5.67) | 10(4.47) |
| **Primary Clinical**<br>**Phenotype (n(%))** |  |  |  |  |  |
| Clinically Normal | 50(100%) | 0(0%) | 22(88%) | 0(0%) | 0(0%) |
| Behavioral Variant<br>FTD | 0(0%) | 8(32%) | 0(0%) | 12(92%) | 15(100%) |
| Primary<br>Progressive<br>Aphasia | 0(0%) | 6(24%) | 1(4%) | 1(8%) | 0(0%) |
| Corticobasal<br>Syndrome | 0(0%) | 5(20%) | 0(0%) | 0(0%) | 0(0%) |
| Alzheimer's<br>Disease<br>Dementia | 0(0%) | 1(4%) | 0(0%) | 0(0%) | 0(0%) |
| MCI – behavior | 0(0%) | 3(12%) | 1(4%) | 0(0%) | 0(0%) |
| MCI - cognitive<br>variants | 0(0%) | 1(4%) | 1(4%) | 0(0%) | 0(0%) |

### Plasma lipid profiles of FTD subjects exhibited numerous differences from control subjects

We examined whether subjects with all types of FTD exhibited differences in plasma lipids compared with controls. Our lipidomic analyses included typical lipids measured after lipid extraction and separate analyses that focused on the more polar glycosphingolipids. We measured ∼530 distinct lipid species across five major lipids classes (free fatty acids, glycerolipids, glycerophospholipids, sphingolipids, and sterols), covering various lipid types (Supplementary Table 1). A Sparse Partial Least Squares–Discriminant Analysis revealed a partial distinction between the FTD and control groups (Figure 2A). As expected, the most abundant lipids in plasma were cholesterol esters (CE), triglycerides (TG), phosphatidylcholine (PC), and sphingomyelin (SM) (Figure 2B-D), which are all components of plasma lipoproteins [19].

**Figure 2.**
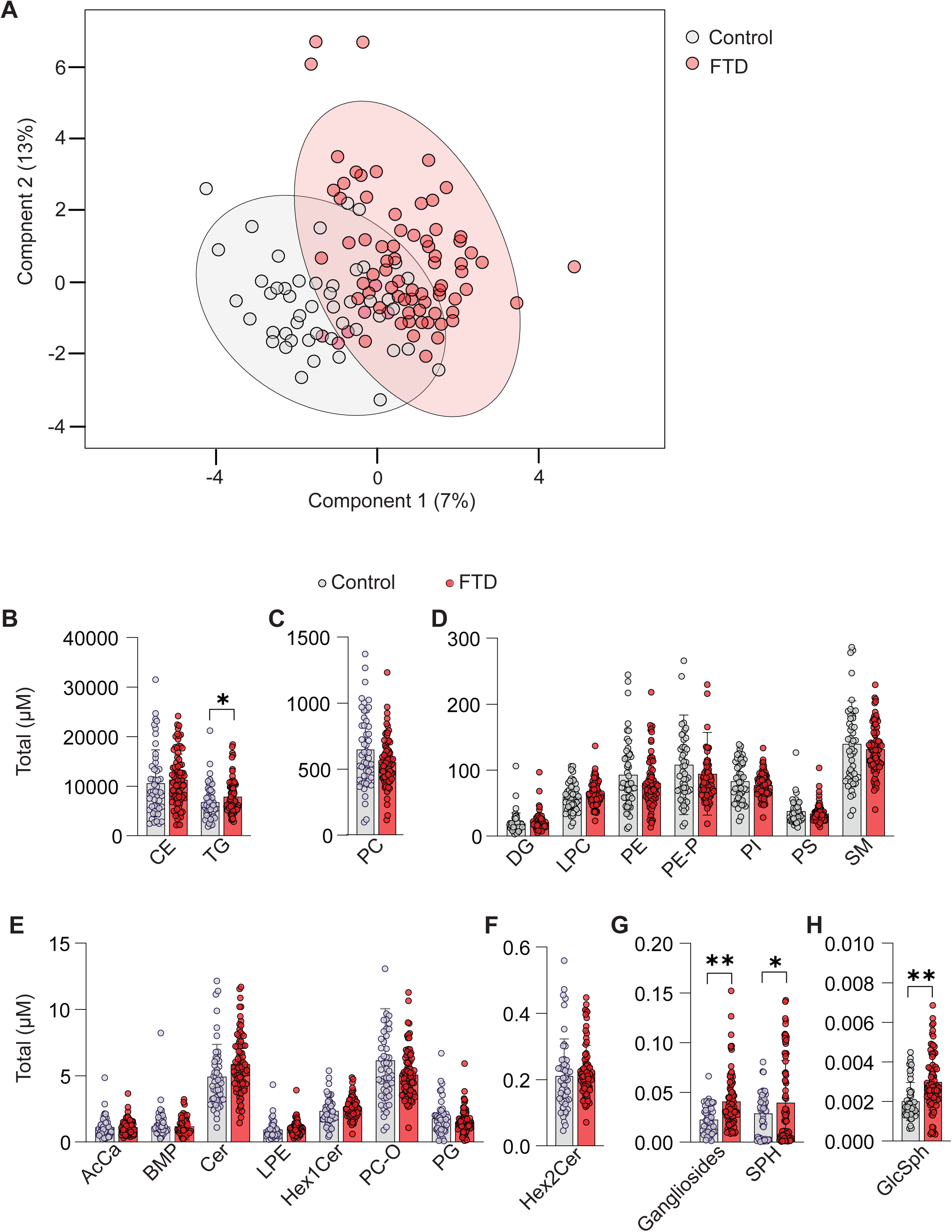
Altered plasma lipid classes of FTD subjects, compared with controls. (**A**) The Sparse Partial Least Squares–Discriminant Analysis revealed a partial distinction between the FTD and control groups. **(B)** The rations of plasma lipid class levels to total lipid levels were compared between FTD and control groups across various lipid classes. Significant alterations were observed in TG, gangliosides, SPH, and GlcSph levels in FTD cases, compared to controls. Data are presented as mean ± standard deviation (mean ± SD). Statistical significance was determined using a parameteric multiple Welch t-test, with *p < 0.05, **p < 0.01.

Plasma levels of TG and several relatively less abundant sphingolipids, including GlcSph, sphingosine (Sph), and gangliosides, were higher in FTD subjects than controls (Figure 2B, G & H). The individual lipid species alterations that are significantly different between FTD subjects and controls are shown in Figure 3A. Additionally, the relationship between the change and average concentration of lipids are shown in Figure 3B. TG species were among the abundant lipids that were increased, as were several low-abundance species of ceramides and GM3.

**Figure 3.**
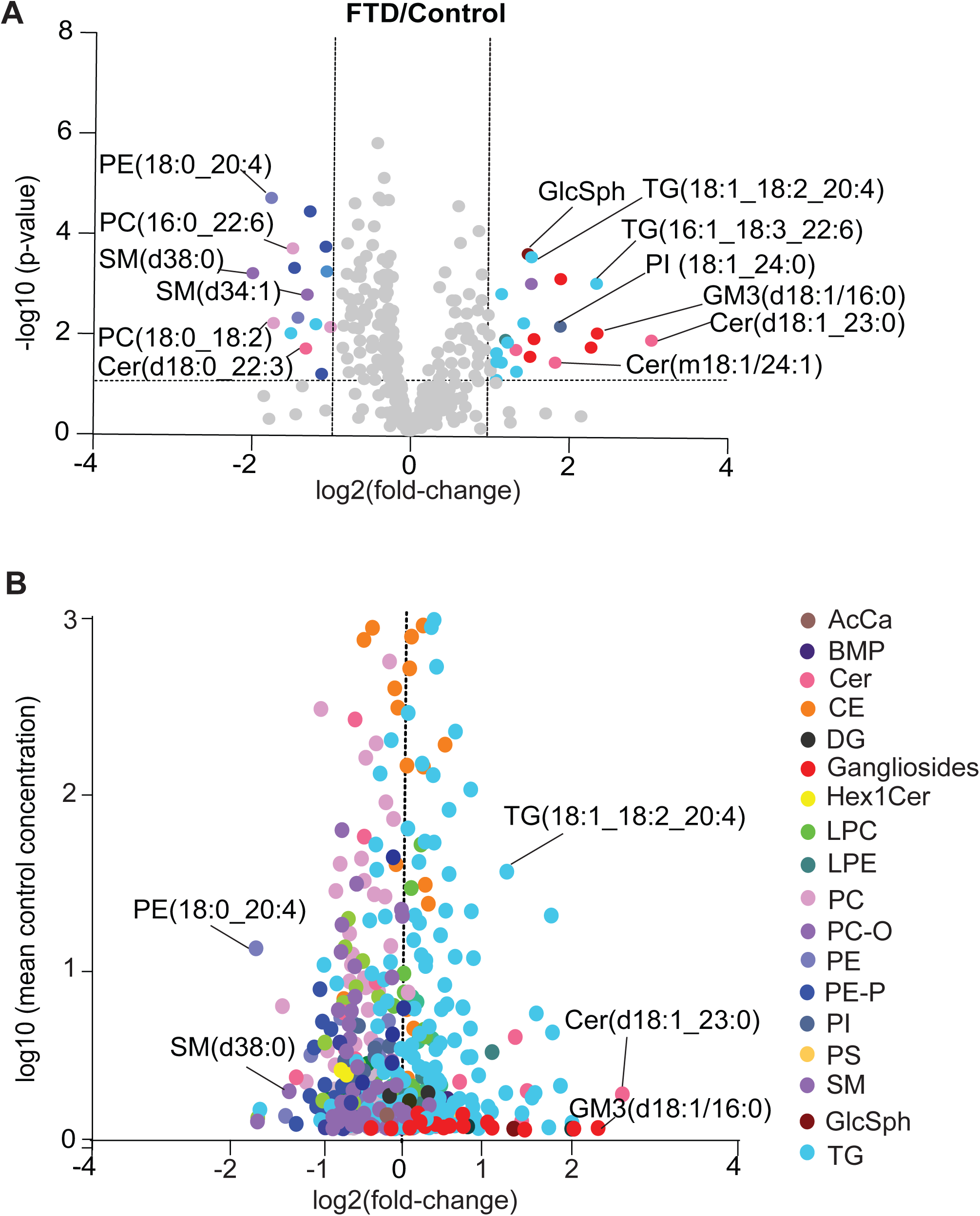
Differential plasma lipid species profiles in FTD subjects, compared with controls. (**A**) Volcano plot illustrating the lipid species with significant fold-changes in plasma levels between FTD cases and controls. Each point represents a lipid, and significant changes are highlighted in different colors (p < 0.05). p values calculated in two sample t-tests. (**B**) Volcano plot illustrating fold-changes for FTD subjects plotted against abundance of lipid species.

Within neutral lipids, plasma TG levels were increased in FTD subjects, and the most elevated TG species included TG with long-chain polyunsaturated fatty acids (e.g., TG(16:0_16:0_20:4), TG(16:1_18:2_20:4), TG(18:1_18:1_22:6), TG(18:1_18:2_20:4) and TG(18:1_18:1_22:5)) (Figure 3A & B and Supplementary Figure 1C).

Levels of plasma PC and PE trended slightly lower in FTD cases than controls. Among the most reduced species were phospholipids with PUFA moieties, such as PC(16:0_22:6), PC(18:2_18:2), and PC(18:0_20:4). Also, some PE species with PUFA were reduced (e.g., PE(18:0_20:4), PE(16:0_22:5), and PE(18:0_18:2)) (Figure 3A &B & and Supplementary Figure 2D).

Among sphingolipids, levels of GlcSph (d18:1) and total gangliosides were 43% (p<0.001) and 49% (P<0.001) increased in FTD cases, respectively, and sphingosine (Sph) levels also tended to be increased (31.7%; p = 0.045) (Figure 2G & H). Among gangliosides, primarily GM3(d18:1_16:0) was elevated in the FTD group compared with controls (134%; p<0.001) (Figure 3A, and Supplementary Figure 1A). Additionally, some ceramides exhibited differences between control and FTD groups. Notably, ceramide species with d18:1 sphingosine, Cer(d18:1_23:0) (Figure 3B, and Supplementary Figure 1B) as well as deoxyceramide Cer(m18:1_24:1), were significantly elevated in FTD, while levels of dihydroxyceramides containing d18:0, such as Cer(d18:0_22:0), were decreased. Sphingomyelins were reduced FTD plasma, particularly SM(d34:1), SM(d38:0), and SM(d40:4) (Figure 3B, Supplementary Table 1).

### Changes in plasma levels of specific lipids correlated with different genetic causes of FTD

We analyzed plasma lipidomes to test for changes that were specific for different genetic causes of FTD. We performed an unsupervised cluster (heatmap) analysis to explore lipid clustering of lipid changes among the FTD study groups (Figure 4). As expected, many species among the 150 lipids that were most changed clustered primarily with other lipids of their class. Among various genetic causes of FTD, plasma samples from FTD-*GRN* exhibited greater increases in specific lipids than other FTD causes, relative to the control samples. This includes significant increases in TGs and various sphingolipids, such as gangliosides and hexosylceramides. The heat map also shows that FTD-*GRN* and FTD-*MAPT* exhibited similar overall changes in phospholipids.

**Figure 4.**
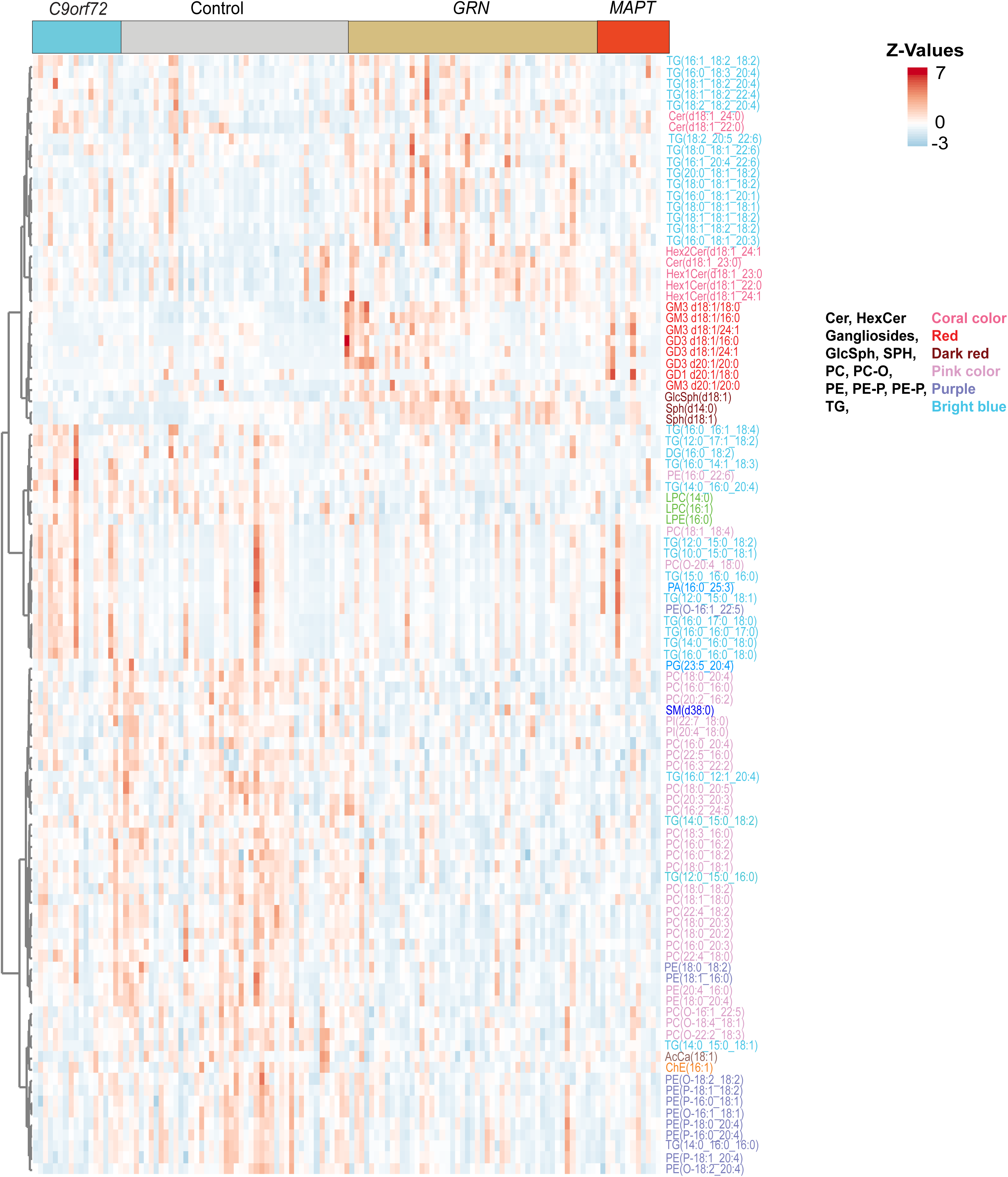
Unsupervised heatmap displaying the top 150 lipid species with the greatest magnitude of change across control, FTD-*GRN*, FTD-*MAPT*, and FTD-*C9orf72* groups. Lipid classes include ceramides (dark red), gangliosides (golden yellow), phospholipids (sky blue, dark green), and triacylglycerols (red). Z-score normalization highlights distinct lipid alterations associated with different conditions.

The data for different genetic forms of FTD were also visualized as volcano plots (Figure 5A-C) and bar graphs for selected lipid species (Figure 5D-F). For FTD-*GRN* patients, sphingolipids such as GlcSph(d18:1), and species of ceramides Cer(d18:1_23:0) and gangliosides (e.g., GM3(d18:1_16:0), GM3(d18:1_18:0), GM3(d18:1_24:1), GM3(d20:1_20:0) and GD3(d18:1_24:1)) were increased. In addition, sphingosine (27%; p=0.07) and hexosylceramides, HexCer(d18:1_24:1) and HexCer (d18:1_23:0), were increased (25%; p=0.09 and 28%; p=0.1, respectively) in FTD-*GRN* subjects (Supplementary Table 1). The GM3(d18:1/16:0) level was also elevated in plasma of FTD-*MAPT* and FTD-*C9orf72* patients (Figure 5B, C & F). GM3(d18:1_18:0) was also elevated in FTD-*MAPT* (Figure 5C &F). However, SM (38:0) was reduced in FTD-*GRN* and FTD-*MAPT*, and SM (34:1) was reduced only in *C9orf72* carriers. TG species, such as TG(18:1_18:2_20:4), were elevated in FTD-*GRN* subjects, and TG(16:0_12:0_20:4) in FTD-*C9orf72* patients, whereas TG(16:0_18:1_20:3) was decreased in MAPT (Figure 5A-C).

**Figure 5.**
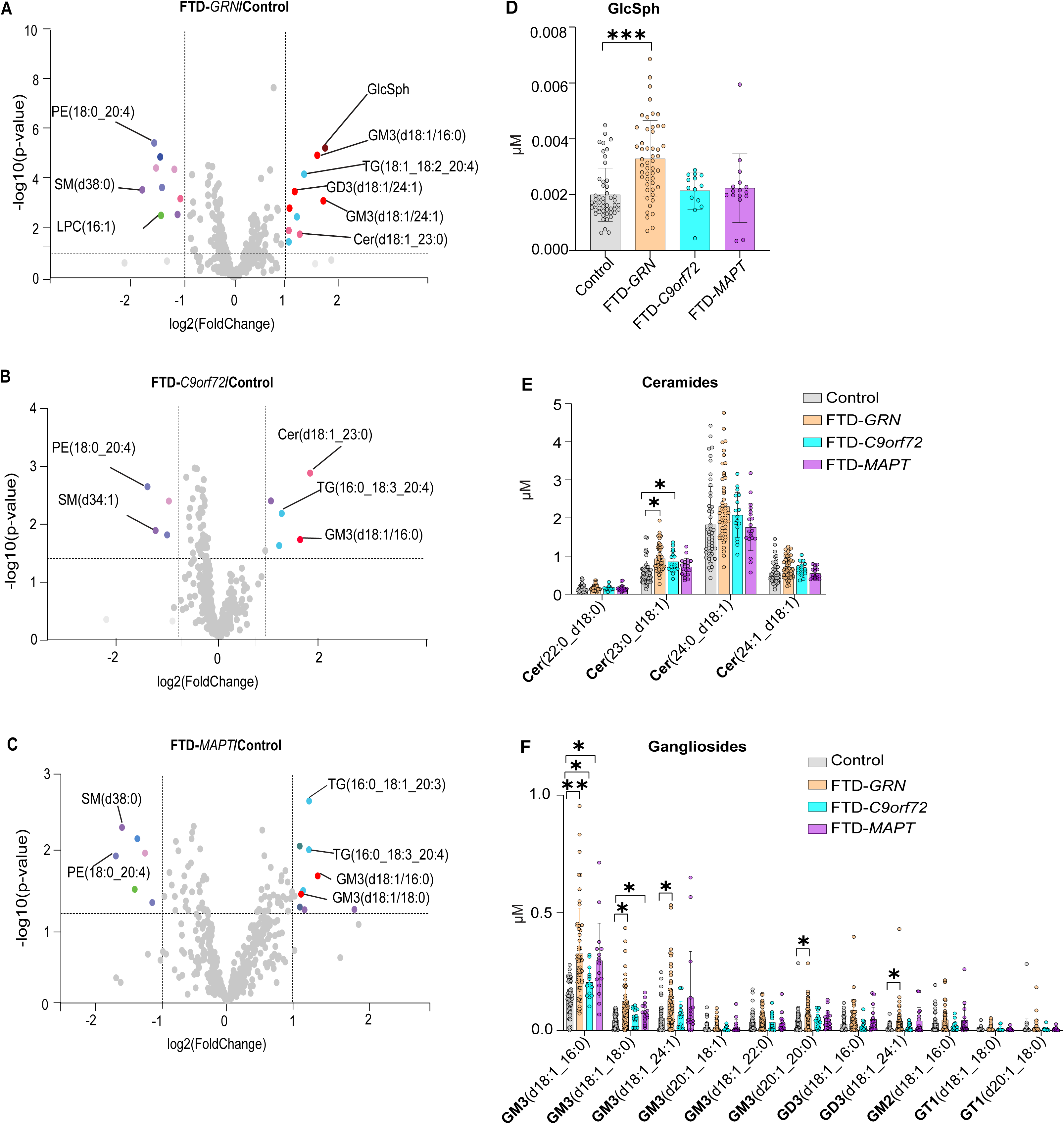
Alterations in the plasma lipid profile of FTD patients with distinct genetic mutations, compared with control subjects. (**A-C**) Volcano plots of differential lipid levels in FTD subtypes, compared to control subjects. These volcano plots present the log2 fold-change (x-axis) and-log10(p-value) (y-axis) of lipid species in plasma from three (FTD) genetic subtypes (FTD-*GRN*, FTD-*C9orf72*, and FTD-*MAPT*), compared to controls. Each point represents a lipid, and the direction of change indicates its relative abundance in FTD patients versus controls. p value calculated in two sample t-tests. **(D)** Plasma GlcSph levels are significantly great in FTD-*GRN* mutations than controls, whereas FTD-*C9orf72* and FTD-*MAPT* subtypes showed no significant differences. **(E)** Specific ceramide species (e.g., Cer (23:0_d18:1)) display distinct alterations, with levels much greated in FTD-*GRN* and FTD-*C9orf72* than controls. **(F)** Ganglioside profiles reveal significant elevation certain species in FTD patients with different genetic mutations (FTD-*GRN*, FTD-*C9orf72*, FTD-*MAPT*). Data are presented as mean ± SD. *p < 0.05, **p < 0.01, ***p < 0.001.

The levels of some phospholipids, such as PC(18:0_22:6) and PC(16:1_22:3), were reduced in FTD-*GRN*, and PC(18:1_18:0) was decreased in FTD-*MAPT*. One species of PE(18:0_20:4) was reduced across FTD patients with different genetic variants. Furthermore, LPC(16:1) levels were reduced in FTD-*GRN* and FTD-*MAPT* (Figure 5A-C).

### Symptomatic FTD-*GRN* subjects exhibited higher levels in plasma glucosylsphingosine, ganglioside GM3, and ceramide species than in asymptomatic subjects

Gangliosides and other sphingolipids are altered in post-mortem brain samples of FTD-*GRN* subjects [4]. Additionally, FTD-*GRN* subjects exhibited the most robust plasma lipid changes in the current study. Therefore, we compared plasma lipids further in symptomatic and asymptomatic FTD-*GRN* patients (Figure 6A &B).

**Figure 6.**
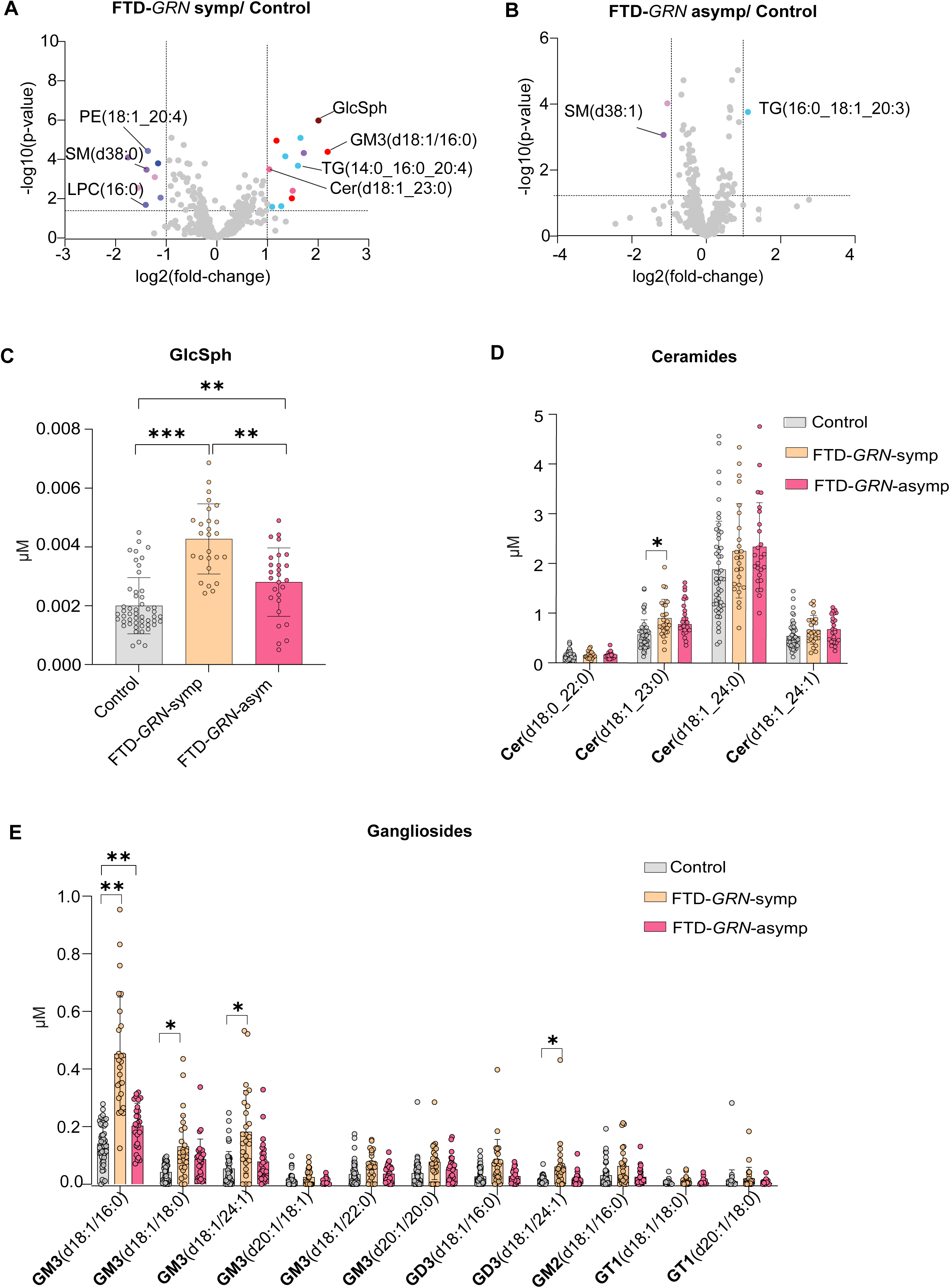
Lipid level differences between symptomatic and asymptomatic FTD-*GRN* subjects and controls. (**A & B**) Volcano plots showing differential lipid levels between FTD-*GRN* symptomatic and symptomatic individuals with controls. Significant lipid species are highlighted. (**C**) Plasma level of GlcSph level in FTD-sym, FTD-asym and controls. **(D)** Plasma levels of ceramide species in FTD-sym, FTD-asym and controls. (**E**) Plasma levels of gangliosides species in FTD-sym, FTD-asym and controls. *p < 0.05, **p < 0.01, ***p < 0.001.

Several lipid species were significantly different in symptomatic FTD-*GRN* subjects relative to controls. For example, GlcSph(d18:1), Cer(d18:1_23:0), GM3(d18:1_16:0), GM3(d18:1_24:1), GM2(d18:1_16:0) and GD3(d18:1_24:1) (Figures 6A-E) were 75%; p<0.001, 38%; p<0.01, 48%; p<0.001, 34%; p<0.01, 32%; p<0.01 and 25%, p<0.05, respectively, higher in symptomatic individuals than controls. In addition, GlcSph(d18:1), GM3(d18:1_16:0), GM3(d18:1_24:1), GM2(d18:1_16:0) and GD3(d18:1_24:1) were 37%; p<0.01, 71%; p<0.01, 54%; p<0.01, 68%; p<0.01 and 32%, p<0.05, respectively, higher in symptomatic individuals than asymptomatic FTD-*GRN* subjects. TG(14:0_16:0_20:4) was higher, and PE(18:1_20:4), SM(38:0) and LPC(16:0) were lower in symptomatic individuals than controls (Figure 6A).

### Plasma levels of GlcSph and specific species of ganglioside, ceramide, sphingomyelin and triglyceride correlated with disease severity and exhibit potential diagnostic utility

To test whether changes in lipid levels correlate with FTD severity, we analyzed lipid species concentrations in relationship to dementia severity. Disease severity was assessed using the six domains of clinical dementia rate (CDR) plus the behavior/comportment and language domains (CDR plus NACC FTLD).

These analyses revealed that altered levels of certain plasma lipids were associated with more severe FTD symptoms. For FTD-*GRN* subjects, GlcSph(d18:1) showed a significant correlation with clinical severity (R = 0.46 p = 0.021) (Figure 7A & Table 2), suggesting a link between lipid metabolism and FTD progression. Similarly, GM3(d18:1_16:0), Cer(d18:1_23:0), and Cer(d18:1_24:1) were positively correlated with disease severity in FTD-*GRN* individuals (R = 0.362, p = 0.01; R = 0.45, p = 0.02; and R = 0.38, p = 0.054, respectively) (Figure 7B-D). In the FTD-*MAPT* group, TG(16:0_18:1_20:3) showed a positive correlation with disease severity (R = 0.56, p = 0.04), while SM(38:0) exhibited a negative correlation (R =-0.48, p = 0.06 (Figure 7E & F). In the FTD-*C9ORF72* subjects (Table 2), SM(d38:1) and SM(d42:2) negatively correlated with severity level CDR + NACC FTLD-SB.

**Figure 7.**
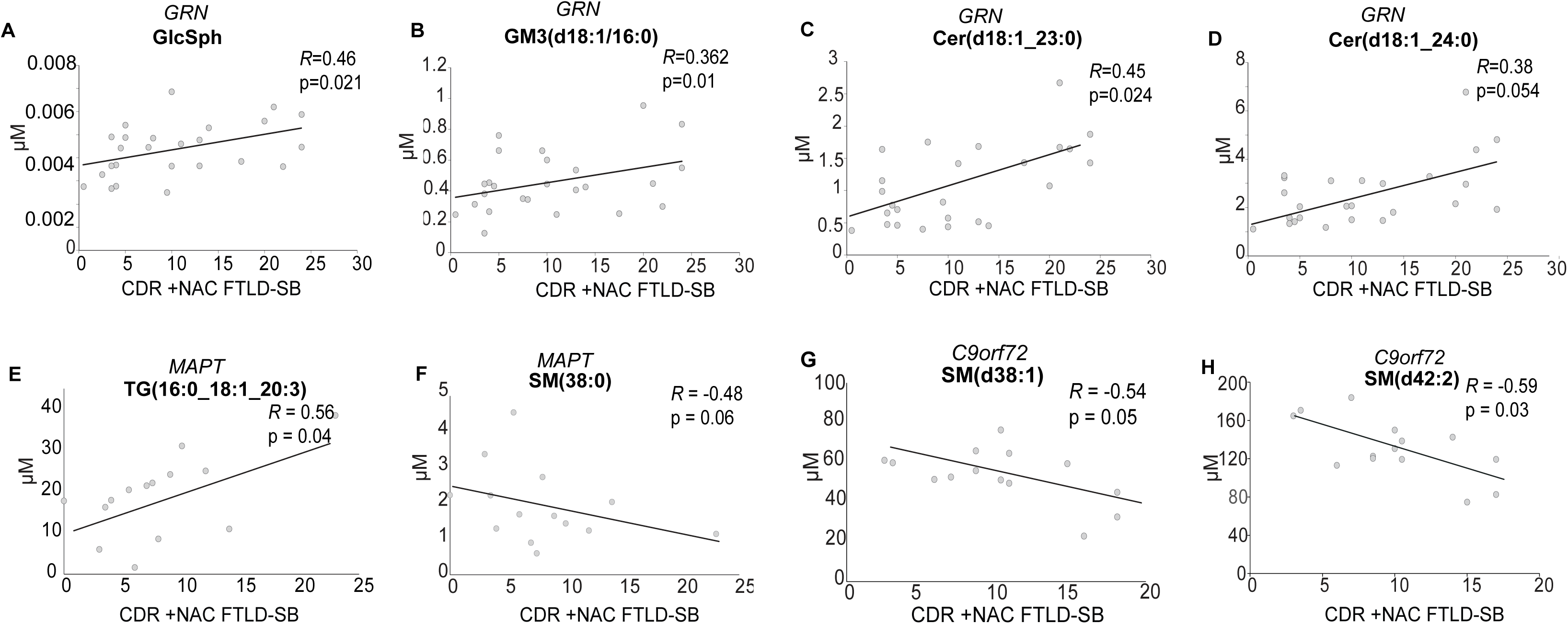
Specific lipid species are associated with disease severity. (**A-D**) Correlation of GlcSph (A), GM3(d18:1/16:0) (B), Cer(d18:1_23:0), and Cer(d18:1_24:0) (D) levels with clinical scores (CDR+NAC and FTLD-SB) in FTD-GRN-sym. **(E-F)** Correlation of TG(16:0_18:1_20:3) and SM(d38:0) (F) levels with clinical scores (CDR+NAC and FTLD-SB) in FTD-*MAPT*-sym. **(G-H)** Correlation of SM(d38:1) (G) and SM(d42:2) (H) levels with clinical scores (CDR+NAC and FTLD-SB) in FTD-*C9orf72*-sym.

**Table 2:**
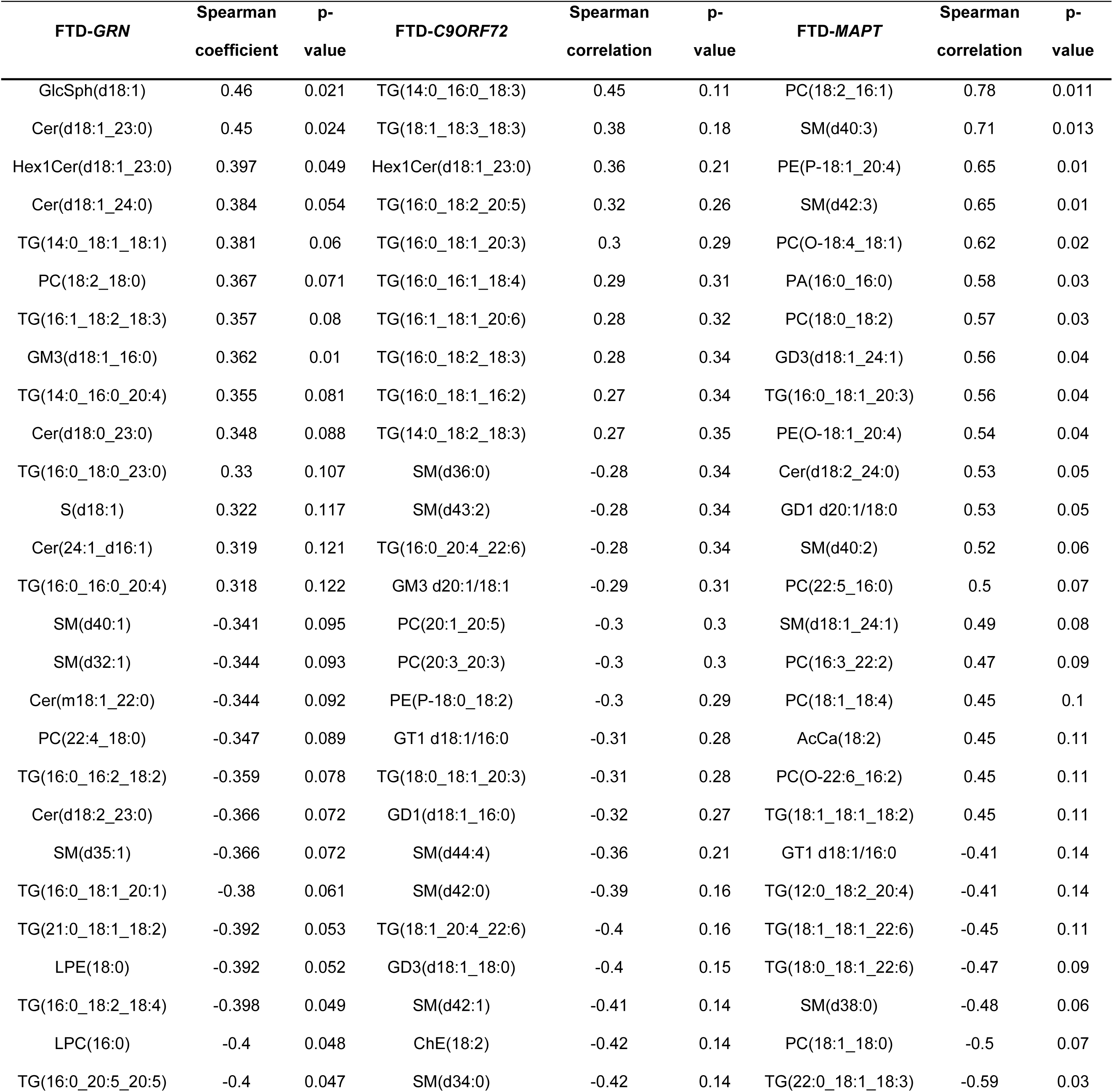
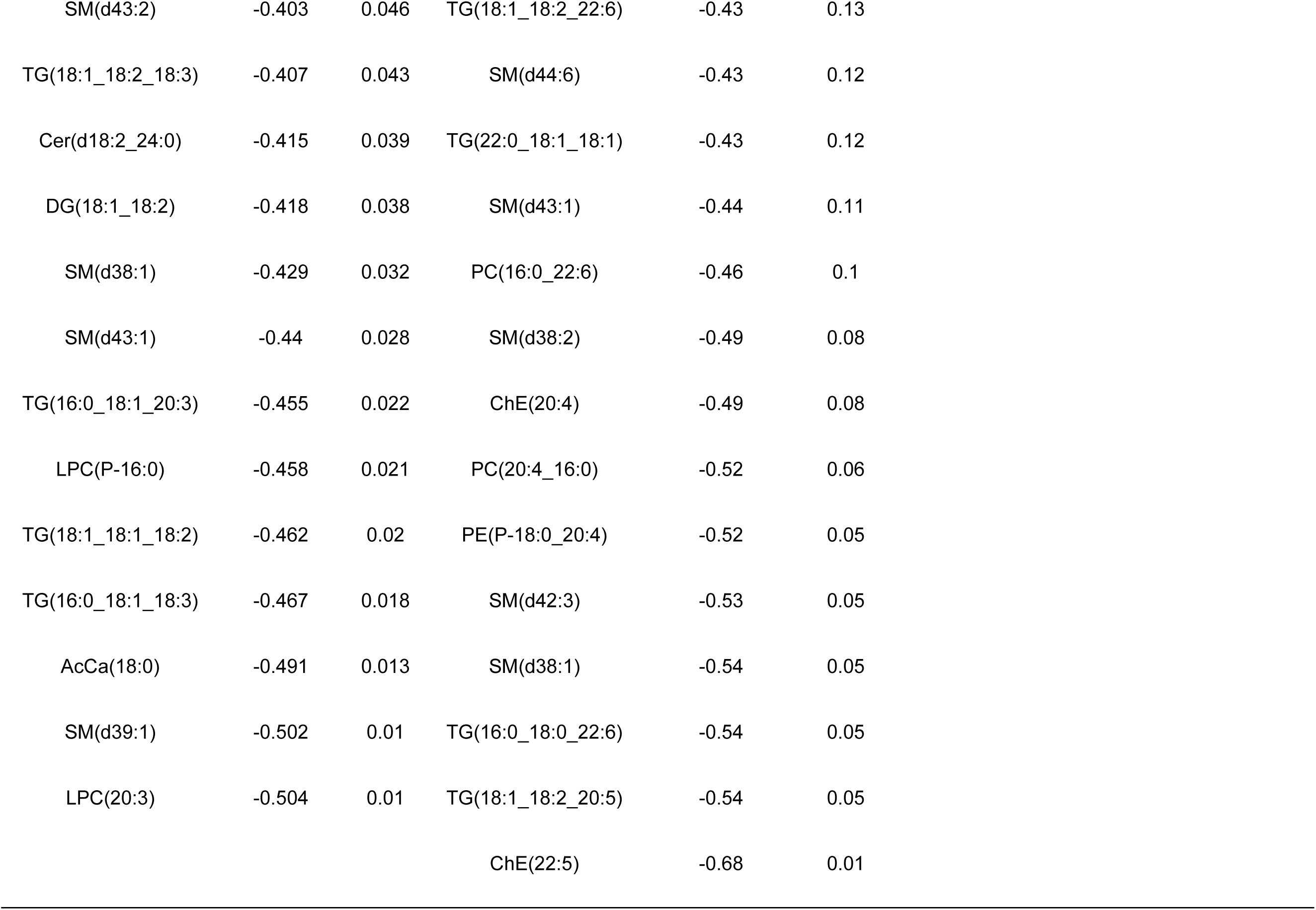
Correlation Between Lipid Species and CDR+NACC FTLD-SB. . Spearman’s correlation coefficients and p-values for lipid species and CDR+NACC FTLD-SB in FTD-GRN, FTD-C9ORF72, and FTD-MAPT patients (CDR+NACC FTLD-SB > 0 at first visit).

| **FTD-GRN** | **Spearman**<br>**coefficient** | **p-**<br>**value** | **FTD-C9ORF72** | **Spearman**<br>**correlation** | **p-**<br>**value** | **FTD-MAPT** | **Spearman**<br>**correlation** | **p-**<br>**value** |
| --- | --- | --- | --- | --- | --- | --- | --- | --- |
| GlcSph(d18:1) | 0.46 | 0.021 | TG(14:0_16:0_18:3) | 0.45 | 0.11 | PC(18:2_16:1) | 0.78 | 0.011 |
| Cer(d18:1_23:0) | 0.45 | 0.024 | TG(18:1_18:3_18:3) | 0.38 | 0.18 | SM(d40:3) | 0.71 | 0.013 |
| Hex1Cer(d18:1_23:0) | 0.397 | 0.049 | Hex1Cer(d18:1_23:0) | 0.36 | 0.21 | PE(P-18:1_20:4) | 0.65 | 0.01 |
| Cer(d18:1_24:0) | 0.384 | 0.054 | TG(16:0_18:2_20:5) | 0.32 | 0.26 | SM(d42:3) | 0.65 | 0.01 |
| TG(14:0_18:1_18:1) | 0.381 | 0.06 | TG(16:0_18:1_20:3) | 0.3 | 0.29 | PC(O-18:4_18:1) | 0.62 | 0.02 |
| PC(18:2_18:0) | 0.367 | 0.071 | TG(16:0_16:1_18:4) | 0.29 | 0.31 | PA(16:0_16:0) | 0.58 | 0.03 |
| TG(16:1_18:2_18:3) | 0.357 | 0.08 | TG(16:1_18:1_20:6) | 0.28 | 0.32 | PC(18:0_18:2) | 0.57 | 0.03 |
| GM3(d18:1_16:0) | 0.362 | 0.01 | TG(16:0_18:2_18:3) | 0.28 | 0.34 | GD3(d18:1_24:1) | 0.56 | 0.04 |
| TG(14:0_16:0_20:4) | 0.355 | 0.081 | TG(16:0_18:1_16:2) | 0.27 | 0.34 | TG(16:0_18:1_20:3) | 0.56 | 0.04 |
| Cer(d18:0_23:0) | 0.348 | 0.088 | TG(14:0_18:2_18:3) | 0.27 | 0.35 | PE(O-18:1_20:4) | 0.54 | 0.04 |
| TG(16:0_18:0_23:0) | 0.33 | 0.107 | SM(d36:0) | -0.28 | 0.34 | Cer(d18:2_24:0) | 0.53 | 0.05 |
| S(d18:1) | 0.322 | 0.117 | SM(d43:2) | -0.28 | 0.34 | GD1 d20:1/18:0 | 0.53 | 0.05 |
| Cer(24:1_d16:1) | 0.319 | 0.121 | TG(16:0_20:4_22:6) | -0.28 | 0.34 | SM(d40:2) | 0.52 | 0.06 |
| TG(16:0_16:0_20:4) | 0.318 | 0.122 | GM3 d20:1/18:1 | -0.29 | 0.31 | PC(22:5_16:0) | 0.5 | 0.07 |
| SM(d40:1) | -0.341 | 0.095 | PC(20:1_20:5) | -0.3 | 0.3 | SM(d18:1_24:1) | 0.49 | 0.08 |
| SM(d32:1) | -0.344 | 0.093 | PC(20:3_20:3) | -0.3 | 0.3 | PC(16:3_22:2) | 0.47 | 0.09 |
| Cer(m18:1_22:0) | -0.344 | 0.092 | PE(P-18:0_18:2) | -0.3 | 0.29 | PC(18:1_18:4) | 0.45 | 0.1 |
| PC(22:4_18:0) | -0.347 | 0.089 | GT1 d18:1/16:0 | -0.31 | 0.28 | AcCa(18:2) | 0.45 | 0.11 |
| TG(16:0_16:2_18:2) | -0.359 | 0.078 | TG(18:0_18:1_20:3) | -0.31 | 0.28 | PC(O-22:6_16:2) | 0.45 | 0.11 |
| Cer(d18:2_23:0) | -0.366 | 0.072 | GD1(d18:1_16:0) | -0.32 | 0.27 | TG(18:1_18:1_18:2) | 0.45 | 0.11 |
| SM(d35:1) | -0.366 | 0.072 | SM(d44:4) | -0.36 | 0.21 | GT1 d18:1/16:0 | -0.41 | 0.14 |
| TG(16:0_18:1_20:1) | -0.38 | 0.061 | SM(d42:0) | -0.39 | 0.16 | TG(12:0_18:2_20:4) | -0.41 | 0.14 |
| TG(21:0_18:1_18:2) | -0.392 | 0.053 | TG(18:1_20:4_22:6) | -0.4 | 0.16 | TG(18:1_18:1_22:6) | -0.45 | 0.11 |
| LPE(18:0) | -0.392 | 0.052 | GD3(d18:1_18:0) | -0.4 | 0.15 | TG(18:0_18:1_22:6) | -0.47 | 0.09 |
| TG(16:0_18:2_18:4) | -0.398 | 0.049 | SM(d42:1) | -0.41 | 0.14 | SM(d38:0) | -0.48 | 0.06 |
| LPC(16:0) | -0.4 | 0.048 | ChE(18:2) | -0.42 | 0.14 | PC(18:1_18:0) | -0.5 | 0.07 |
| TG(16:0_20:5_20:5) | -0.4 | 0.047 | SM(d34:0) | -0.42 | 0.14 | TG(22:0_18:1_18:3) | -0.59 | 0.03 |

|  |  |  |  |  |  |
| --- | --- | --- | --- | --- | --- |
| SM(d43:2) | -0.403 | 0.046 | TG(18:1_18:2_22:6) | -0.43 | 0.13 |
| TG(18:1_18:2_18:3) | -0.407 | 0.043 | SM(d44:6) | -0.43 | 0.12 |
| Cer(d18:2_24:0) | -0.415 | 0.039 | TG(22:0_18:1_18:1) | -0.43 | 0.12 |
| DG(18:1_18:2) | -0.418 | 0.038 | SM(d43:1) | -0.44 | 0.11 |
| SM(d38:1) | -0.429 | 0.032 | PC(16:0_22:6) | -0.46 | 0.1 |
| SM(d43:1) | -0.44 | 0.028 | SM(d38:2) | -0.49 | 0.08 |
| TG(16:0_18:1_20:3) | -0.455 | 0.022 | ChE(20:4) | -0.49 | 0.08 |
| LPC(P-16:0) | -0.458 | 0.021 | PC(20:4_16:0) | -0.52 | 0.06 |
| TG(18:1_18:1_18:2) | -0.462 | 0.02 | PE(P-18:0_20:4) | -0.52 | 0.05 |
| TG(16:0_18:1_18:3) | -0.467 | 0.018 | SM(d42:3) | -0.53 | 0.05 |
| AcCa(18:0) | -0.491 | 0.013 | SM(d38:1) | -0.54 | 0.05 |
| SM(d39:1) | -0.502 | 0.01 | TG(16:0_18:0_22:6) | -0.54 | 0.05 |
| LPC(20:3) | -0.504 | 0.01 | TG(18:1_18:2_20:5) | -0.54 | 0.05 |
|  |  |  | ChE(22:5) | -0.68 | 0.01 |

Receiver operating characteristic (ROC) analysis, performed for FTD-*GRN* cases, showed that the two most significant lipid species, GlcSph(d18:1) and GM3(d18:1_16:0), discriminated moderately well between all FTD-*GRN* cases and controls, with AUC values of 0.782 and 0.814, respectively (Figure 8A). In symptomatic FTD-GRN cases, performance improved, particularly for GM3(d18:1_16:0), with AUCs of 0.848 and 0.925 for GlcSph and GM3, respectively (Figure 8B). By contrast, both markers were less effective in distinguishing asymptomatic carriers from controls (AUCs of 0.708 and 0.716; Figure 8C).

**Figure 8.**
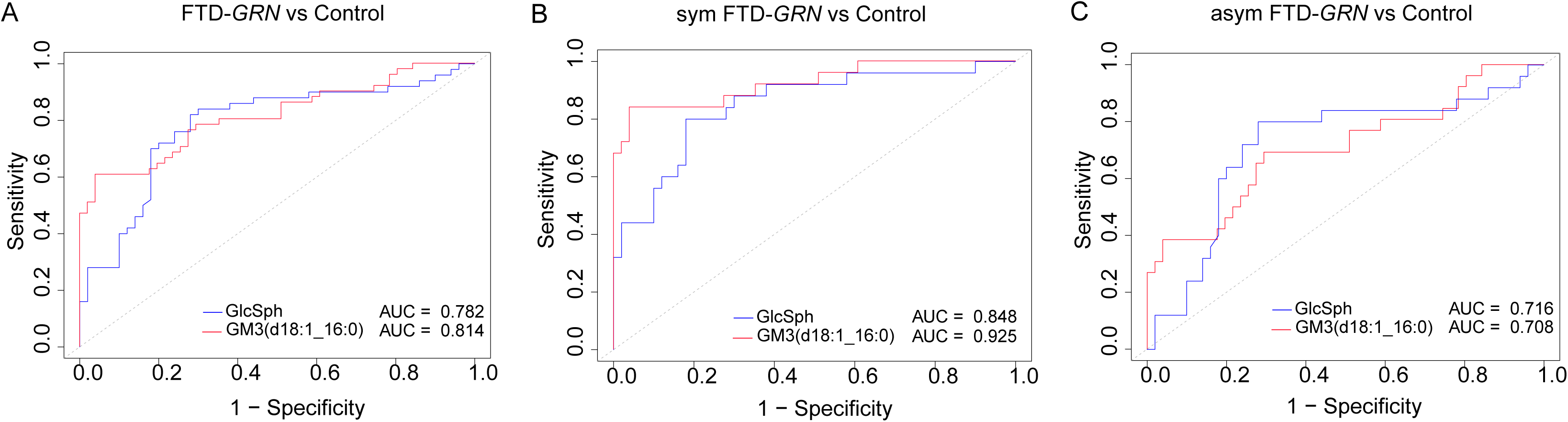
ROC curves illustrating the discriminatory performance of GlcSph(d18:1) and GM3(d18:1_16:0) across all FTD-*GRN*, symptomatic and asymptomatic groups and controls. (**A**) ROC analysis comparing the overall FTD-*GRN* group to controls, with GlcSph(d18:1) and GM3(d18:1_16:0) showing AUC values of 0.782 and 0.814, respectively. (**B**) In the symptomatic FTD-*GRN* subgroup versus controls, GlcSph(d18:1) and GM3(d18:1_16:0) exhibited higher diagnostic performance with AUCs of 0.848 and 0.925, respectively. (**C**) The asymptomatic FTD-*GRN* subgroup showed modest discrimination from controls, with AUCs of 0.708 for GlcSph(d18:1) and 0.716 for GM3(d18:1_16:0). Graphs show AUC values with 95% confidence intervals.

## Discussion

Recent progress studying FTD suggests that at least some forms of the disease are linked to altered lipid metabolism [4, 20–22]. Here, we report that the concentration of numerous lipids, predominantly specific species of sphingolipids and of glycerolipids, were altered in FTD subjects in general. These alterations exhibited both shared and distinct patterns across genetic variants of FTD, and levels of several lipids correlated with disease severity.

One of the least common lipids in plasma, GlcSph is a well-recognized biomarker for disease progression in Gaucher’s disease, where it accumulates due to mutations in the lysosomal enzyme (GCase) that normally degrades this glucosylsphingolipid [23]. We found significant changes in GlcSph levels in FTD subjects (Figure 3B). Subgroup analyses showed that the increased levels in FTD subjects were due to the increase in the FTD-*GRN* subgroup (Figure 5A & D). Consistent with our findings, previous reports showed that plasma GlcSph was increased in FTD-*GRN* mutation carriers, but not in other symptomatic genetic mutation groups or non-carriers [15, 17, 18]. However, we found a greater change in GlcSph levels in FTD-*GRN* plasma samples than was reported.

Whether plasma GlcSph levels reflect solely brain disease is unclear. PGRN deficiency impairs lysosomal function and disrupts sphingolipid metabolism in various cell types, leading to accumulation of several lipids, including GlcSph [21, 24]. Since lysosomal dysfunction due to *GRN* mutation affects many cell types, including circulating leukocytes [23, 25], elevated plasma GlcSph could be derived from blood cells.

Several lysosomal storage disorders result in ganglioside accumulation in the brain [4, 26], and we reported increased levels of gangliosides in FTD-*GRN* post-mortem brains [4]. In this study, we found a marked increase of GM3(d18:1_16:0) in all cases of FTD, and these changes were most pronounced in plasma from symptomatic *GRN* mutation carriers. Altered lipid metabolism may not be limited to FTD due to *GRN* mutations since elevated ganglioside levels were found across patients with different mutations (e.g., GM3(d18:1_16:0), GM3(d18:1_18:0) in FTD-*C9orf72* and FTD-*MAPT*) [27]. Gangliosides were reported to be increased with *GRN-* deficiency due to dramatically reduced levels of lysosomal BMP, a lipid co-factor required for efficient degradation of complex glycolipids [4, 28]. In our study, we detected no significant changes in plasma levels of BMP in FTD subjects, compared with controls. This suggests that plasma BMP levels may not accurately reflect changes in tissues, such as the brain. Notably, reduced urinary levels of BMP in FTD-*GRN* have been reported [29], a finding that requires further investigation.

Among plasma sphingolipids, several ceramide species exhibited different patterns in FTD subjects. For instance, Cer(d18:1_23:0) levels were significantly elevated in plasma of all FTD patients, and Cer(d18:1_24:1) tended to be elevated in FTD, whereas levels of Cer(d18:0_22:0) were reduced. This is consistent with the growing recognition that different ceramides can have different and, sometimes, opposite functions in physiology [30]. The increase in plasma ceramides containing acyl chains with 23 carbons is noteworthy. Odd-numbered acyl chains are rare in humans and are either synthesized from microbial precursors or due to rare start of fatty acid synthesis from propionyl CoA rather than acetyl-CoA. Propionyl-CoA can accumulate with mitochondrial dysfunction, as in FTD and ALS, and is genetically associated with neurological disease. These findings merit further investigation for these ceramides as potential biomarkers. Of note, we also found lower levels of some sphingomyelin species, including SM(d34:1), SM(d38:0), and SM(d40:2), in FTD-*C9orf72*, FTD-*GRN*, FTD-*MAPT,* respectively, suggesting more broadly de-regulated sphingolipid metabolism.

Specific species of TG were elevated across FTD patients, and the most upregulated TG species included TG(18:1_18:2_20:4), TG(16:0_16:0_20:4). Notably, these TGs contained long-chain polyunsaturated fatty acids. TG levels are increased in serum of subjects with bvFTD [31, 32] and in the brains of FTD-*GRN* patients [21]. We lack an explanation for the observed changes in plasma PUFA-TG levels in FTD patients, although this finding suggests a metabolic shift from other glycerolipid species, such as phospholipids, to TG [33].

Some species of phospholipids were lower in FTD cases than controls. Among PC species most decreased in FTD were PC(16:0_22:6), PC(18:2_18:2), and PC(18:0_20:4). PE species, such as PE(18:0_20:4) and PE(16:0_22:5), also exhibited significant decreases. The findings are similar to previous studies that showed the level of phospholipids reduced in FTD group, even though the total level of PS and PG appeared to be reduced in these investigations particularly [31]. The reduction of PUFAs in phospholipid species with a concomitant increase of their levels in TG may indicate a metabolic shift towards and more saturated glycerophospholipids and a compensatory increase of PUFAs in TG lipids.

In our search for biomarkers that correlate with disease progression, changes in gangliosides and other sphingolipid degradation products were most pronounced in symptomatic subjects with FTD-*GRN* mutations, and we found a positive correlation between GlcSph, GM3(d18:1/16:0), Cer(d18:1_23:0), and Cer(d18:1_24:1) levels and FTD severity, as measured by the CDR + NACC FTLD-SB dementia scoring system. SM(d38:1) and SM(d42:2) negatively correlated with severity levels in CDR + NACC FTLD-SB in FTD-*C9ORF72*. Additionally, TG(16:0_18:1_20:3) showed a positive correlation, while SM(d38:0) exhibited a negative correlation with severity level CDR + NACC FTLD-SB in FTD-*MAPT*. This suggests a potential utility of these lipids as biomarkers for tracking disease progression. Similar correlations were reported in AD, where gangliosides were linked to disease severity [17, 18]. In addition, the ROC analysis findings suggest that GlcSph(d18:1) and GM3(d18:1_16:0) possess strong discriminatory capacity, particularly in symptomatic FTD-*GRN* patients, highlighting their potential as lipid-based biomarkers for disease detection and progression monitoring.

These studies suggest including these sphingolipids in biomarker panels for monitoring disease severity.

A limitation of our study is the relatively modest sample sizes within each group despite the multi-center design of the study, reflecting the inherent challenge in the collection of plasma samples from a sizable cohort of different genetic FTD cases in the United States. This constraint may limit the robustness of statistical analyses, particularly in correlation assessments with clinical measures. We also focused on genetic causes of FTD rather than non-genetic, sporadic cases. Moreover, our study subjects were primarily Caucasian in ethnic origin; studies of subjects with FTD from other ethnicities are clearly needed. Finally, this study is cross-sectional, which precludes causal inferences about the role of lipid changes in FTD progression. While we observed significant lipidomic differences between FTD and control subjects, as well as correlations with disease severity, longitudinal studies will be needed to validate these findings, assess changes in lipid levels over time, and confirm whether these alterations are consistently predictive of disease progression.

## Methods

*Plasma samples and participants*. Plasma samples were collected from de-identified patients with FTD and aged-matched controls. These samples included symptomatic carriers of mutations in GRN (sym-GRN, n=25), symptomatic GRN carriers (asym-GRN, n=25) C9orf72 (C9orf72, n=15), and MAPT (n=15), and aged-matched controls (n=50) Figure 1A. Samples were obtained from the National Centralized Repository for Alzheimer’s Disease and Related Dementias and the University of California, San Francisco. Clinical diagnoses, based on comprehensive evaluation, were established by a behavioral neurologist at the time of assessment [34, 35]. Demographic and clinical characteristics are detailed in Table 1.

*Lipidomics extraction.* Plasma was diluted (1+9, v:v) with 150 mM aqueous ammonium hydrogen carbonate to improve sample handling and accuracy. A variant of the MTBE (methyl tert-butyl ether)-methanol-water lipid extraction method [36] was performed in 1.5-mL polypropylene Eppendorf safe-lock tubes. One hundred microliters of 1:10 diluted plasma (corresponding to 10 µL plasma) were combined with 1 mL of MTBE/methanol (7+2, v:v) containing internal standards (10 µL of SPLASH II LIPIDOMIX Mass Spec Standard and 100 pmol of Cer(d18:1-d7/15:0) and 100 µL of deionized water. To ensure the quality and precision of the results, pooled quality control samples were included for every 20 study samples. Samples were agitated on an orbital shaker (Eppendorf Thermomixer C; 30 min, 1,000 rpm, 4°C), and centrifuged in an Eppendorf table-top centrifuge (10,000 g, 10 min, 4°C). The upper lipid-containing phase was transferred to a new safe-lock tube and evaporated in a vacuum centrifuge. The dried lipid film was reconstituted in 100 µL of 65:30:5 (isopropanol:acetonitrile:water) solution for lipidomics and BMP lipid analysis. All liquid transfers were performed with an Opentron OT2 system (Opentrons Labworks, Inc).

*Glucosylsphingosine extraction.* The GlcSph extraction method was adapted from a published protocol [15]. Briefly, 75 μL of plasma was mixed with 200 μL of a methanol-based internal standard mixture containing 5 ng/mL of GlcSph-d5 (Avanti Polar Lipids cat # 860636P-1mg) in polypropylene tubes. The mixture was vortexed for 1 min, allowed to stand at room temperature for 15 min, and then centrifuged at 10,000 g for 10 min. The resulting supernatant was collected and directly injected into the mass spectrometry (MS) system for analysis.

*Ganglioside extraction*. Lipids were extracted from 100 µL of plasma using 1 mL of methanol in 1000 rpm, 4 °C for 2 h. The extraction mixture was centrifuged at 10,000 rpm for 10 min to pellet proteins, and the solvent layers containing gangliosides and other metabolites were collected in an Eppendorf tube and dried under speed vac. Dried extract was reconstituted in 1 mL of LC-MS-grade water and desalted by Sola HRP SPE 30 mg/2 mL 96-well plate (Thermo Scientific #60509-001). Desalting cartridges were cleaned three times with 1 mL of methanol, equilibrated three times with LC-MS-grade water, and then the extracts dissolved in water were loaded onto the cartridge and washed three times with water, and finally, gangliosides were eluted by 3 mL of methanol. Eluates were dried under N_2_ and reconstituted in chloroform:methanol:water (120:60:9).

*Lipidomics.* The HPLC-MS method was adopted from [37]. Briefly, the HPLC conditions in this experiment involved Dionex UltiMate 3000 HPLC system equipped with a Waters C18 (2.1 × 150 mm, 1.7 μm) liquid chromatography column for the separation of lipid mixtures. The temperature of the sample tray and column were maintained at 4 and 55 °C, respectively. The mobile phase composition for liquid chromatography consisted of mobile phase A (10 mM ammonium formate in 40:60 water:acetonitrile + 0.1% formic acid) and mobile phase B (10 mM ammonium formate in 90:10:isopropanol:acetonitrile:water + 0.1% formic acid). The gradient (v:v) used was as follows: 0–1.5 min, held at 30%; 1.5–4 min, linear gradient from 32% to 45% B; 4–5 min, linear gradient from 45% to 52% B; 5–8 min, linear gradient from 52% to 58% B; 8–11 min, linear gradient from 58% to 66% B; 11–14 min, linear gradient from 66% to 70% B; 14–18 min, linear gradient from 70% to 75% B; 18–21 min, linear gradient from 75% to 97% B; 21–25 min, held at 97%; 25–25.1 min, returned to 30% B; and 25.1–30 min, equilibrated at 30% B. The flow rate of the mobile phase was set at 0.3 mL/min, and the injection volume was 2 μL.

The MS analysis was performed using a Q-Exactive Orbitrap (QE) MS (Thermo Fisher Scientific, Waltham, MA) equipped with an electrospray ionization (ESI) source. The instrument parameters were as follows. The spray voltage was set to 4.2 kV, and the heated capillary and the HESI were held at 320 and 300 °C, respectively. The S-lens RF level was set to 50, and the sheath and auxiliary gases were set to 35 and 3 units, respectively. These conditions were held constant for both positive and negative ionization mode acquisitions. External mass calibration was performed using the standard calibration mixture every 7 days.

MS spectra of lipids were acquired in the full-scan/data-dependent MS2 mode. For the full-scan acquisition, the resolution was set to 70,000, the AGC target was 1e6, the maximum injection time was 50 msec, and the scan range was *m*/*z* = 133.4–2000. For data-dependent MS2, the top 10 ions in each full scan were isolated with a 1.0-Da window, fragmented at a stepped normalized collision energy of 15, 25, and 35 units, and analyzed at a resolution of 17,500 with an AGC target of 2e5 and a maximum injection time of 100 msec. The underfill ratio was set to 0. The selection of the top 10 ions was subject to isotopic exclusion with a dynamic exclusion window of 5.0 sec. Processing of raw data was performed using LipidSearch 5.0 software (Thermo Fisher Scientific/Mitsui Knowledge Industries). Further filtering and normalization were conducted using an in-house developed, web-based app, *LipidCruncher*, which will be reported elsewhere. Semi-targeted quantification was performed by normalizing the area under the curve (AUC) to the AUC of internal standards.

*Glucosylsphingosine analysis.* For each sample, 10 µL of extracted sample was injected on a HALO HILIC 2.0 µm, 3.0 × 150 mm column (Advanced Materials Technology, PN 91813-701) using a flow rate of 0.45 mL/min at 45°C. Mobile phase A consisted of 92.5/5/2.5 ACN/IPA/H2O with 5 mM ammonium formate and 0.5% formic acid. Mobile phase B consisted of 92.5/5/2.5 H2O/IPA/ACN with 5 mM ammonium formate and 0.5% formic acid. The gradient was programmed as follows: 0.0–2 min at 100% B, 2.1 min at 95% B, 4.5 min at 85% B, hold to 6.0 min at 85% B, drop to 0% B at 6.1 min and hold to 8.5 min. For the analysis GlcSph, Vanquish Horizon UHPLC system coupled to OE240 Exactive Orbitrap MS (Thermo Fisher Scientific) equipped with a heated electrospray ionization probe.

MS settings included an ion transfer tube temperature of 300°C, vaporizer temperature of 275°C, Orbitrap resolution of 120,000 for MS1 and 30,000 for MS2, RF lens at 70%, with a maximum injection time of 50 ms for MS1 and 54 ms for MS2. Positive and negative ion voltages were set at 3250 and 2500 V, respectively. Gas flow rates included auxiliary gas at 10 units, sheath gas at 40 units, and sweep gas at 1 unit. High-energy collision dissociation (HCD) fragmentation was stepped at 15, 25, and 35%, and data-dependent tandem MS (ddMS2) ran with a cycle time of 1.5 s, an isolation window of 1 m/z, an intensity threshold of 1.0e4, and a dynamic exclusion time of 2.5 s.

Full-scan mode with ddMS^2^ was performed over an m/z range of 250-700, with EASYICTM used for internal calibration. The raw data were processed and aligned with LipidSearch 5.0, using a precursor tolerance of 5 ppm and a product tolerance of 8 ppm.

*Analysis of bis(monoacylglycero)phosphate (BMP).* Lipids were separated using ultra-high-performance liquid chromatography (UHPLC) coupled with tandem mass spectrometry (MS/MS). UHPLC analysis was conducted on a C30 reverse-phase column (Thermo Acclaim C30, 2.1 x 150 mm, 2.6 μm) maintained at 50°C and connected to a Vanquish Horizon UHPLC system, along with an OE240 Exactive Orbitrap MS (Thermo Fisher Scientific) equipped with a heated electrospray ionization probe. Each sample (5 μL) was analyzed in both positive and negative ionization modes. The mobile phase included 60:40 water:acetonitrile with 10 mM ammonium formate and 0.1% formic acid, while mobile phase B consisted of 90:10 isopropanol:acetonitrile with the same additives. The chromatographic gradient involved: Initial isocratic elution at 30% B from-3 to 0 minutes, followed by a linear increase to 43% B (0-2 min), then 55% B (2-2.1 min), 65% B (2.1-12 min), 85% B (12-18 min), and 100% B (18-20 min). Holding at 100% B from 20-25 min, a linear decrease to 30% B by 25.1 min, and holding from 25.1-28 min. Flow rate of 0.26 mL/min, injection volume of 2 μL, and column temperature of 55°C. Mass spectrometer settings included an ion transfer tube temperature of 300°C, vaporizer temperature of 275°C, Orbitrap resolution of 120,000 for MS1 and 30,000 for MS2, RF lens at 70%, with a maximum injection time of 50 ms for MS1 and 54 ms for MS2. Positive and negative ion voltages were set at 3250 V and 2500 V, respectively. Gas flow rates included auxiliary gas at 10 units, sheath gas at 40 units, and sweep gas at 1 unit. High-energy collision dissociation (HCD) fragmentation was stepped at 15%, 25%, and 35%, and data-dependent tandem MS (ddMS2) ran with a cycle time of 1.5 s, an isolation window of 1 m/z, an intensity threshold of 1.0e4, and a dynamic exclusion time of 2.5 s.

Full-scan mode with ddMS^2^ was performed over an m/z range of 250-1700, with EASYICTM used for internal calibration. The raw data were processed and aligned with LipidSearch 5.0, using a precursor tolerance of 5 ppm and a product tolerance of 8 ppm. Further filtering and normalization were conducted using an in-house app, Lipidcruncher. Semi-targeted quantification was performed by normalizing the area under the curve (AUC) to the AUC of internal standards. Samples for BMP analysis were run in both negative and positive ion mode. All the quantification presented in the figures of the manuscript come from the negative-mode analysis. Positive-mode analysis of BMP was used only to confirm the identify of BMP species.

*Analysis of gangliosides*. The HILIC-MS method was adopted from [38] to separate and detect gangliosides. HPLC analysis was performed with a Phenomenex (Thermo Fisher Scientific, CAT#, 2.0 × 150 mm, operated at 60 °C; Bremen, Germany). 15 μL of each sample was injected and acquired in negative ionization mode. The mobile phase A consisted of acetonitrile with 0.2% formic acid and mobile phase B consisted of 10 mM aqueous ammonium acetate, pH 6.1, adjusted with formic acid. Column equilibration was performed using 12.3% B for 5 min prior to each run. Chromatographic condition: mobile-phase gradient as follows: 0 min: 87.7% A + 12.3% B; and 15 min: 77.9% A + 22.1% B. The re-equilibration time between runs was 5 mins. The flow rate for the separation was set to 0.6 mL/min. The column oven temperature was set to 40 °C, and the temperature of the autosampler tray was set to 4 °C. The spray voltage was set to −4.5 kV, and the heated capillary and the HESI were held at 300 °C and 250 °C, respectively. The S-lens RF level was set to 50, and the sheath and auxiliary gas were set to 40 and 5 units, respectively. These conditions were held constant during acquisition.

External mass calibration was performed using the standard calibration mixture every 7 days. MS spectra of lipids were acquired in full-scan/data-dependent MS2 mode. For the full-scan acquisition, the resolution was set to 70,000, the AGC target was 1e6, the maximum injection time was 50 msec, and the scan range was *m*/*z* = 700–2500 in the negative ion mode. For data-dependent MS2, the top 10 ions in each full scan were isolated with a 1.0-Da window, fragmented at stepped normalized collision energies of 25, 35, and 50 units, and analyzed at a resolution of 17,500 with an AGC target of 2e5 and a maximum injection time of 100 msec. The underfill ratio was set to 0. The selection of the top 10 ions was subject to isotopic exclusion with a dynamic exclusion window of 5.0 sec. Processing of raw data was performed in Xcalibur™ software (Thermo Fisher Scientific).

*Statistical analyses.* Analyses were performed using R (version 4.4.1, GNU General Public License v2), GraphPad Prism 10, Metaboanlytes 6.0 and illustrator software. Plasma lipidomics were compared between the studied groups and the mutation types using either Student’s *t* test (2-tailed), Mann-Whitney-Wilcoxon test or parameteric multiple Welch t-test and 1-way or 2-way ANOVA for multiple comparisons. Due to missing age or sex information for certain subjects in the selected population, no demographic adjustments were made in the analysis. Unsupervised heatmap analysis of the top 150 changes in lipid species was conducted using Metaboanalytes 6.0 [39]. To examine the association between lipid levels and baseline level in CDR+NACC FTLD-SB score in sym-*GRN, c9orf72 and MAPT* subjects, Spearman’s coefficient correlation was used. Spearman’s rank correlation, a non-parametric statistical method, was used to estimate the strength and direction of association without assuming a linear relationship between biomarkers and CDR+NACC FTLD-SB scores. This analysis includes only the initial available sample in the dataset that corresponds to a CDR+NACC FTLD-SB score greater than zero for each subject. Asymptomatic groups were defined as the carriers with CDR+NACC FTLD-SB scores consistently equal to 0. Symptomatic groups were defined as carriers with CDR+NACC FTLD-SB scores consistently exceeding 0. For all analyses, *P*-values < 0.05 were considered statistically significant. Receiver operating characteristics (ROC) curve analyses were plotted, and the area under the curve (AUC) including 95% confidence interval (CI) values are reported.

*Study approval.* All human plasma samples were collected in accordance to ethical guidelines and protocols. Plasma biospecimens and data were provided by the ARTFL & LEFFTDS Consortia (ARTFL: U54 NS092089, funded by the National Institute of Neurological Diseases and Stroke and National Center for Advancing Translational Sciences; LEFFTDS: U01 AG045390, funded by the National Institute on Aging and the National Institute of Neurological Diseases and Stroke. This study was approved by the local institutional review boards for all ARTFL and LEFFTDS clinical sites. Informed consent was obtained from all participants.

## Acknowledgments

We thank members of Farese & Walther for helpful discussions. This work was supported by a grant from the Bluefield Project to Cure FTD (to R.V.F. and T.C.W.), and postdoctoral fellowship grants from the Bluefield Project to Cure FTD (to Y.A. and S.S.). T.C.W. is a Howard Hughes Medical Institute Investigator. We acknowledge support from an NIH/NCI Cancer Center Support Grant (Core grant P30 CA008748) to MSKCC.

## Author contributions

Y.A.A., P.A.L., A.L.D., R.V.F., and T.C.W. conceived the project, and R.V.F. and T.C.W. acquired project funding. P.A.L. and A.L.D., provided plasma samples. Y.A.A. generated reagents, performed lipidomics experiments and data analysis. P.A.L. and A.L.D. provided clinical data. S.B. generated preliminary data and discussed data. A.H. and S.S. helped with scientific discussion and data analysis. Y.A.A., R.V.F., and T.C.W. co-wrote the manuscript with input from all authors.

---

**Supplementary Figure 1.**
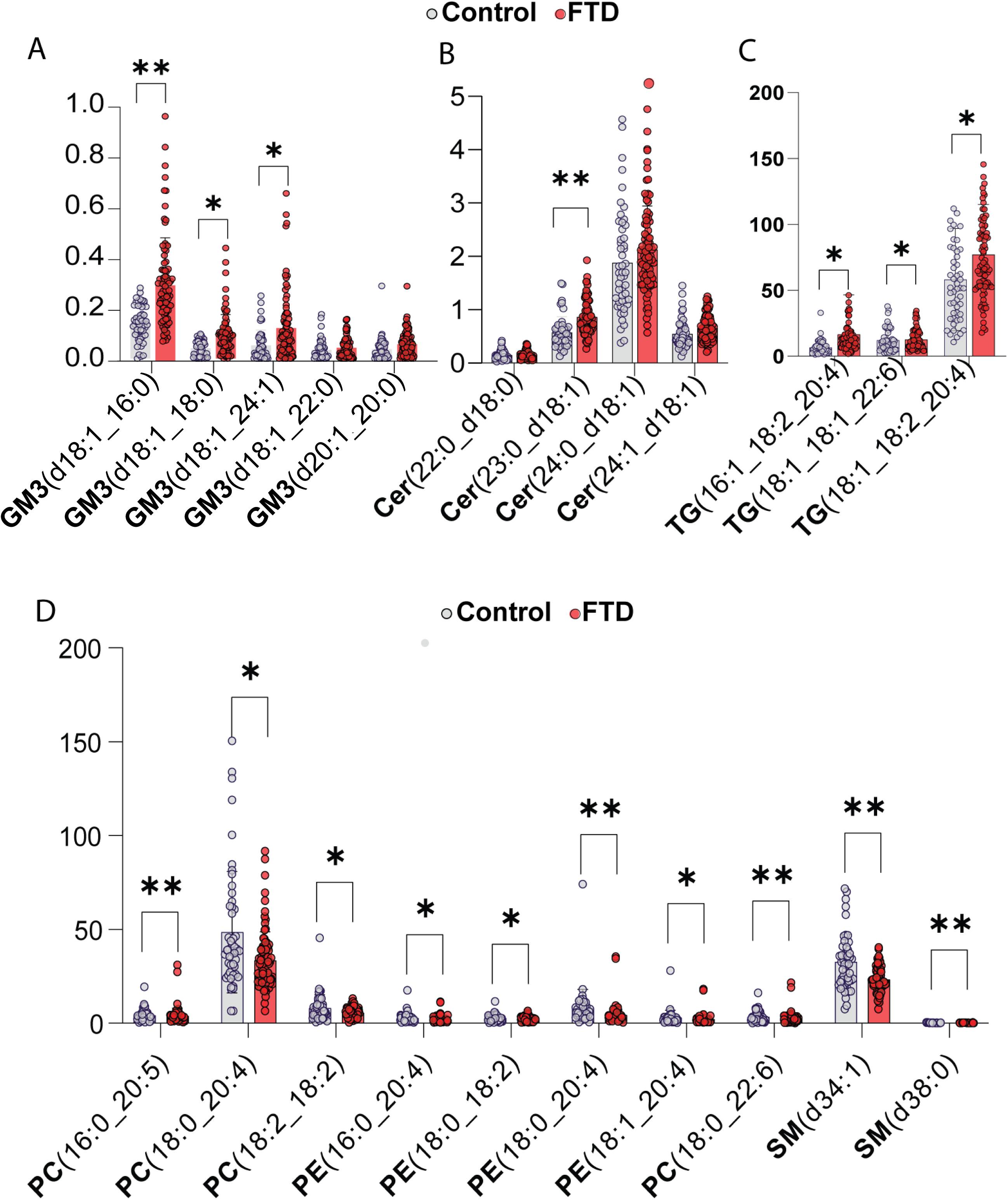
Alteration of plasma lipid species in FTD, compared to controls. (**A-C**) Levels of specific plasma ganglioside species (A), certain ceramide species (B), and certain TG species were significantly higher in FTD cases than controls. (**D**) Levels of specific plasma phospholipid and sphingomyelin species were lower in FTD cases than controls. Bars represent the mean ± standard deviation (SD). Each dot represents to samples. Data are presented as mean ± SD, *p<0.05, **p<0.01 (multiple parametric tests, with Welch t-test).

